# Acyl-AcpB, a FabT co-repressor in *Streptococcus pyogenes*

**DOI:** 10.1101/2023.06.09.544320

**Authors:** Clara Lambert, Alice d’Orfani, Marine Gaillard, Qiufen Zhang, Karine Gloux, Claire Poyart, Agnès Fouet

**Author notes:** Address correspondence to Agnes Fouet, and Clara Lambert. Department of Molecular Biology, Umeå University, Umeå, Sweden (Clara Lambert). Medicinal Chemistry and Bioinformatics Center, Shanghai Jiao Tong University School of Medicine, Shanghai China.

## Abstract

Membranes are a universal barrier to all cells. Phospholipids, essential bacterial membrane components, are composed of a polar head and apolar fatty acid (FA) chains. Most bacterial FA are synthesized by the FA synthesis pathway (FASII). In streptococcaceae, enterococci and *Lactococcus lactis*, a unique feedback mechanism controls the FASII gene expression. FabT, encoded in the FASII main locus, is the repressor and it is activated by acyl-ACP. Many Streptococci, *Enterococcus faecalis*, but not *L. lactis*, possess two ACPS. AcpA encoding gene is within the FASII locus and is, to a certain extent, coregulated with the FASII genes. Acyl-AcpA is the end product of FASII. AcpB encoding gene is in operon with *plsX*. The role of AcpB as FabT corepressor is controversial. *Streptococcus pyogenes*, which causes a wide variety of diseases ranging from mild non-invasive to severe invasive infections, possesses AcpB. In this study, we show by comparing gene repression of FASII genes in wild-type, *fabT* mutant and *acpB* mutant strains grown in the presence and the absence of exogenous FAs, that AcpB is *S. pyogenes* FabT main co-repressor. Also, *acpB* deletion impacts the membrane FA composition and adhesion to eucaryotic cells, highlighting the role of AcpB.

## Introduction

Cell membranes commonly comprise a phospholipid bilayer composed of a polar head and an apolar body constituted of fatty acids (FA). FA structure and length are decisive for membrane topology and properties such as fluidity, permeability and integrity. These features are crucial for the adaptation of bacteria to various environments (1). The FAs are synthesized by the FA synthesis pathway (FASII) that is widespread among bacteria. It consists of an initiation phase followed by a recursive elongation cycle, which produces acyl-acyl carrier protein (acyl-ACP). Unsaturated FAs are produced *via* a shunt of this cycle that is catalyzed by FabM in streptococci. In contrast to the FASII pathway, the regulation of the FASII genes is not conserved. In streptococci, enterococci and *Lactococcus lactis*, a unique feedback mechanism controls the expression of the FASII genes, involving a repressor, FabT, encoded in the FASII main locus. Depending on the *Streptococcus* species, the *fabM* gene is either not controlled or less repressed by FabT than the *fabK* gene, inducing, in addition to lengthening of the FA chains, a shift in the proportion of saturated – unsaturated FAs in *fabT* mutant strains (2, 3).

FabT, a member of the MarR family of regulators, possesses an unconventional corepressor, acyl-ACP [(4) for review (5)]. FabT – acyl-ACP binding to its DNA target varies according to the acyl length, increasing with long chain acyls (4). It also depends on the saturation state of the FA, the *cis* unsaturated FA maximizing FabT-DNA affinity (6). The role of FabT as a FASII repressor in the presence of exogenous FAs (eFAs) can readily be observed by the repression of the FASII genes in the wild-type strain grown in the presence of eFA and the loss of this repression in *fabT* mutant strains grown similarly (6, 7).

In addition to acyl moiety variation, the ACP may differ. Indeed, in many streptococci, as well as in *Enterococcus faecalis*, but not in *L. lactis*, there are two ACPs. AcpA is encoded within the FASII locus; the AcpB encoding gene is found in the same operon as *plsX*, PlsX is involved in the phospholipid biosynthetic pathway (8-10). Whereas *acpA* expression is, to a limited extent with repression folds weaker than those of most other FASII genes, controlled by FabT, that of *acpB* is not (6, 7). The role of AcpB in FASII gene control is controversial. FabT – acyl-ACP DNA binding affinity tests using *S. pneumoniae* purified FabT and acyl-AcpA or acyl-AcpB indicated that only AcpB increased it (8). In contrast, in *E. faecalis*, no difference was observed in the *fabT* gene expression between wild-type and *acpB* mutant strains, grown either in the presence or absence of eFAs (6).

*Streptococcus pyogenes* possesses two ACPs. *S. pyogenes* is a Gram positive strictly human pathogen that yields a wide range of clinical manifestations from mild superficial infections to more life-threatening invasive infections including necrotizing fasciitis (NF), streptococcal toxic shock syndrome (STSS). GAS infections are also responsible for post-infectious complications (11). Altogether, GAS infections are responsible for approximately 517,000 deaths annually worldwide (12). We have previously shown that a mutation in *S. pyogenes fabT* has consequences in membrane FA composition, eFA incorporation, bacterial adhesion and growth in the presence of eukaryotic cells (7).

To determine whether, and to what extent, AcpB is a FabT corepressor in *S. pyogenes*, we studied the effects of deleting *acpB* on FASII gene expression in bacteria grown in the presence or absence of eFAs. We also determined the membrane FA composition in these bacteria as well as the consequences of *acpB* deletion on bacteria - host cells interaction.

## Results and discussion

### *S. pyogenes* AcpB is a FabT corepressor

A strain expressing a FabT^H105Y^ point mutation, mFabT, was previously constructed in the reference strain M28PF1 (an *emm28* strain, referred to as wild-type; WT) (10)Lambert, 2022 #3;Lambert, 2023 #17}. An *acpB* deleted strain, ∧AcpB, was constructed in the WT strain; as a control for the ΔAcpB strain, we used the BWT strain that was obtained during the ΔAcpB strain construction; its *acpB* gene is wild-type and the ΔAcpB and BWT strains share a single surreptitious mutation (Table S1). Growths of WT, mFabT, ΔAcpB and BWT strains were compared in the laboratory media THY and in THY supplemented with an exogenous unsaturated FA source (here 0.1 % Tween 80) (Fig. S1). All strains grew similarly.

To analyze the role of AcpB as a FabT co-repressor, the effects of AcpB on the expression of a set of genes, containing FASII genes and the coregulated *degV*_*1638*_ gene, were determined either in the absence or in the presence of Tween 80 and compared to those of the WT and the mFabT strains (Fig. 1) (5, 7). qRT-PCR were performed on RNA extracted from six conditions: WT, ΔAcpB and mFabT strains, each grown in THY and THY-Tween. To allow the direct comparison of the gene expression in all three strains, the expression of the set of genes was first compared in the BWT and the WT strains and found to be unchanged (Fig. S2). Data were then analyzed by comparing results pairwise between ΔAcpB and WT or mFabT and WT strains in each medium (Fig. 1A-B). In THY, the expression of *fabK* is differentially expressed, in fact overexpressed, in both mutant strains (Fig. 1A). In THY-Tween, all genes were more expressed in both mutant strains than in the WT strain. (Fig. 1B). These data suggested that AcpB is involved in the FASII gene repression in the absence and in the presence of eFAs.

**Figure 1.**
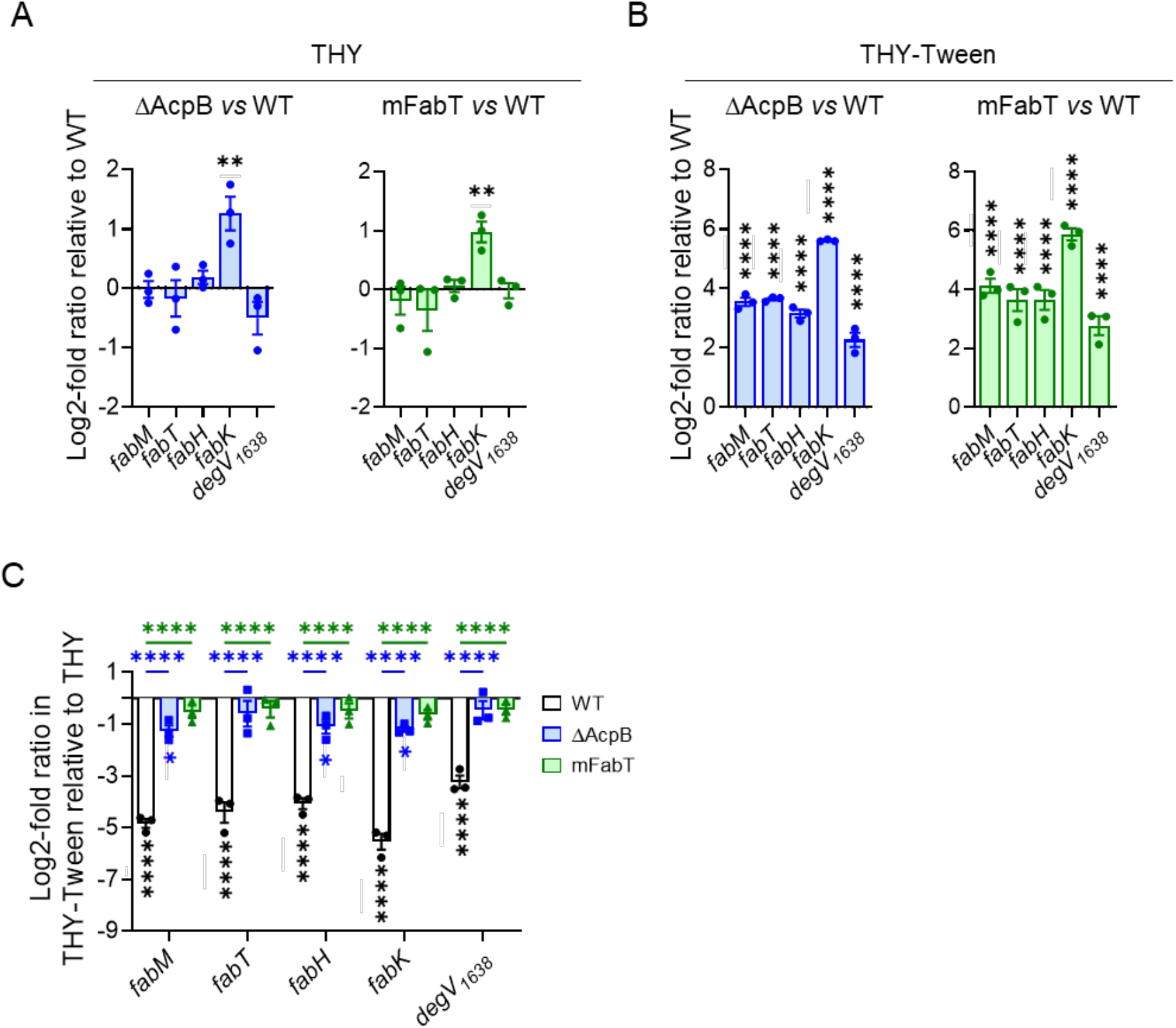
The ΔAcpB strain has lost most of the eFA-related transcriptional repression capacity. WT, ΔAcpB and mFabT strains were grown in THY (A, C) or THY-Tween (B, C) and RNAs were quantified by qRT-PCR. Expression was normalized to that of *gyrA*; relative gene expression is expressed as the log2-fold ratio in ΔAcpB *versus* WT strain (right, A and B) and mFabT *versus* WT strain (left, A and B). C) The relative gene expression is expressed as the log2-fold ratio in a given strain grown in THY-Tween *versus* in THY. White, WT; blue, ΔAcpB, green mFabT. Significance of differences: stars below bars, in the two media for a given strain; stars above bars, in the two media between WT and ΔAcpB or WT and mFabT. No statistically significant diffrences were found between ΔAcpB and mFabT. N =3. 2-way ANOVA, Bonferroni post-test, *, p<0.05; **, p < 0.01; ****, p<0.0001.

To compare the repression defect in the presence of eFAs in the mFabT and ΔAcpB strains, we analyzed the repression exerted in each strain in the presence of Tween 80 (Fig. 1C, stars below the columns). Whereas this addition led to a repression fold between 12 and 25 in the WT strain, it was just above 2 for three out of the five genes in the ΔAcpB strain, and below 2 for all genes in the mFabT strain. This indicates that, in contrats to the mFabT strain, the ΔAcpB strain has not lost all repression activity. We then compared whether the repression exerted in each strain was different from that in the other strains (Fig. 1C, stars above the columns). That in the WT strain is different from that of both mutant strains. The variation of the expression ratio between the ΔAcpB and the mFabT was not statistically different. Altogether, these results confirm that *S. pyogenes* AcpB is a major player in the transcriptional repression of the FASII genes, but that FabT can exert part of its repression activity in the absence of AcpB.

Our data are in sharp contrast with those obtained in *E. faecalis*, when FASII gene expression was assessed in an *acpB* mutant strain, but in agreement with the conclusion drawn from results obtained by a biochemical approach in *S. pneumoniae* (6, 8). Hence, our data strongly suggest that in *S. pyogenes* AcpB is FabT main corepressor and that this may be the case in many *Streptococcus* species. Furthermore, AcpB would exert its role both in the absence and the presence of eFAs. Altogether this indicates that AcpB role may be different in various species and warrants further analysis in other organisms.

### AcpB deletion modifies the *S. pyogenes* membrane FA composition

Since the AcpB deletion leads to a deregulation of FASII gene expression, and the mFabT strain produced longer chain saturated FA and had a decreased unsaturated: saturated FA ratio we tested the consequences of the deletion of *acpB* on FA composition when bacteria were grown in the absence or presence of Tween 80 (Fig. 2A-B). We compared pairwise the membrane FA composition of the ΔAcpB strain to, on the one hand, that of the WT strain and, on, the other hand, of the mFabT strain. As a control, the membrane FA composition of the BWT and the WT strains was compared and found to be similar (Fig. S3, Table S2).

**Figure 2.**
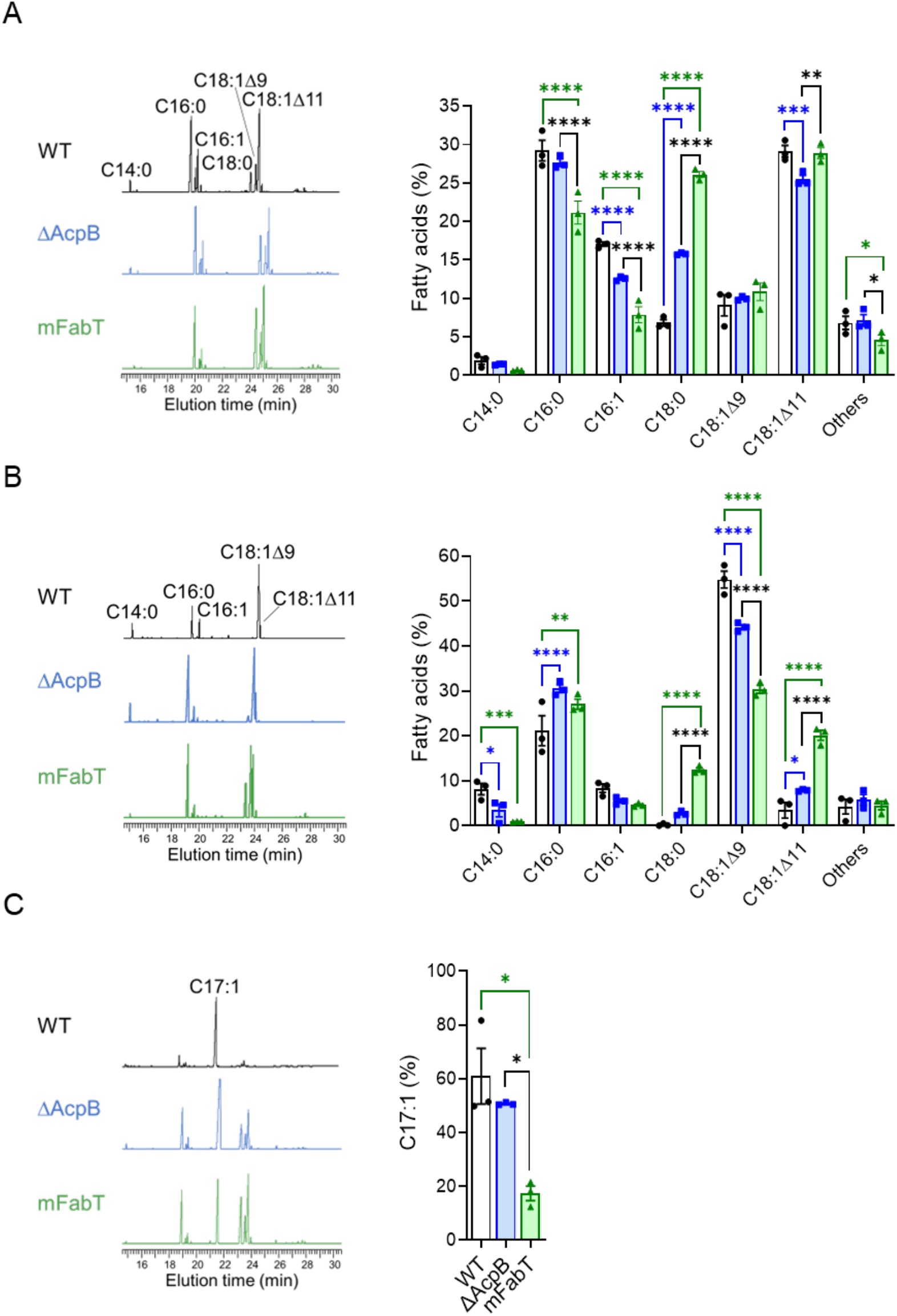
The FA membrane composition of the ΔAcpB strain differs from that of WT and mFabT strains. FA composition of WT, ΔAcpB and mFabT strains grown in A) THY medium B) THY-Tween; C) THY-C17:1. A, B, C) left, FA profiles; right, quantified proportions of major FAs. N=3 (Table S2). A, B) right, 2way ANOVA, Bonferroni post-test, C) one-way ANOVA, Bonferroni post-test; *, p<0.05 **, p<0.01; ***, p<0.001; ****, p<0.0001.

The striking differences, in the absence of Tween 80 between the WT and the ΔAcpB membrane FA composition, were that the ΔAcpB membrane contained less unsaturated FA (C16:1 and C18:1Δ11) and more saturated FA (C18:0) than the WT strain (Fig. 2A). The higher amount of C18:0 in the ΔAcpB (15.78 % *vs* 6.82 %) altered the FA membrane composition in favor of more saturated and longer chain FA. These differences are of the same sort as those found with the mFabT strain as also previously described (Fig. 2A) (7). However, the ΔAcpB membrane contained more C16:1 (27.64 % *vs* 21.15 %) and less C18:0 (15.78 % *vs* 26.08 %) than that of the mFabT strain. In the presence of Tween 80, the ΔAcpB membrane contained 1.2-fold less C18:1Δ9 (44.18 % *vs* 54.8 %) and two-fold more C18:1Δ11 (7.88 % *vs* 3.41 %) than the WT strain (Fig. 2B). Again, these modifications were more important in the mFabT than in the ΔAcpB strain (two-fold less C18:1Δ9 and six-fold more C18:1Δ11 compared to the WT strain). Altogether, the differences observed in the absence or the presence of eFAs between the ΔAcpB and the WT strains are not as important as those between the mFabT and the WT strains. As mentioned, repression was totally lost in the mFabT strain but not in the ΔAcpB strain (Fig. 1C, stars below the columns). This difference may yield a threshold of enzyme synthesis that is amplified at the level of FA synthesis.

The weaker C18:1Δ9 percentage in the ΔAcpB membrane than in that of the WT strain suggested an incorporation defect, as observed with the mFabT strain (Fig. 2B) (7). However, this was not the case; we evaluated FA incorporation in medium supplemented with C17:1 (THY-C17:1), which is not synthesized by bacteria (Fig. 2C). The percentage of C17:1 was similar in WT and ΔAcpB strains grown in THY-C17:1 supplemented medium, and different from that in the mFabT strain (Fig. 2C). The larger C18:1Δ9 decrease observed in the mFabT than in the ΔAcpB strain is due to this incorporation defect difference (Fig. 2C). This difference between *fabT* and *acpB* mutant strains had already been described in *E. faecalis* (6).We previously demonstrated that the absence of FASII gene repression is responsible for the incorporation defect in the mFabT strain. The residual regulation observed in the ΔAcpB strain thus enables incorporation, and the eFAs incorporation difference is a consequence of the regulation difference between the mFabT and ΔAcpB strains (Fig. 1C, stars below the columns).

Our data indicate that AcpB, as a FabT corepressor, controls the membrane FA composition. They also support that AcpB is the main but not the sole FabT co-repressor. This further suggests that in the absence of AcpB, acyl-AcpA is also a corepressor. Moreover, since the mFabT and AcpB membrane FA composition is different both in the presence and the absence of eFAs, acyl-AcpA may operate in both conditions; acyl-AcpA results from the interaction of *de novo* synthesized acyls and in the presence of eFAs, from the reverse PlsX reaction that catalyzes the production of Acyl-ACP from acyl-phosphate (5).

### The ΔAcpB strain displays, a defect in adhesion to eucaryotic cells defect

The mFabT strain displays defects in its adhesion capacity to eucaryotic cells and its growth capacity in cell-conditioned supernatant (7). The ΔAcpB strain membrane FA composition was in-between that of the WT and the mFabT strains. We thus wondered whether the ΔAcpB strain would display the same defects. As for all previous phenotypes, the BWT strain had the same adhesion and growth capacities as the WT strain (Fig. S4). The ΔAcpB strain displayed, like the mFabT strain, an adhesion defect (Fig. 3A). This indicates that the weak regulation produced by the ΔAcpB strain and its consequently intermediate membrane FA composition did not impact the adhesion defect. On the contrary, variations from the wild-type strain are sufficient to impact adhesion.

**Figure 3.**
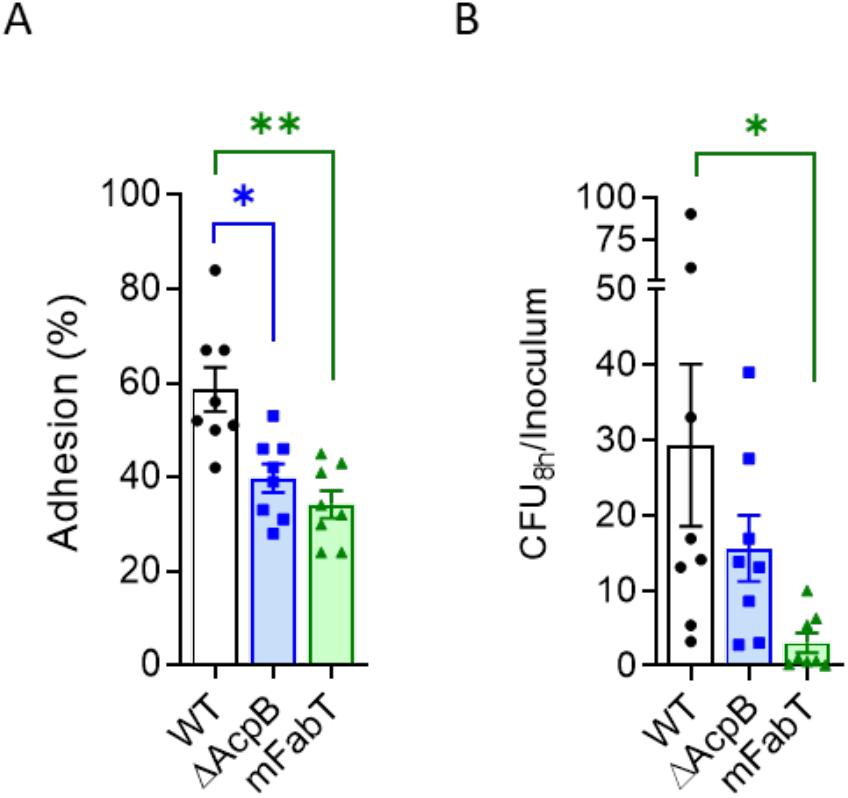
The ΔAcpB mutant strain has an adhesion defect to human cells. Adhesion to human cells and growth capacities in cell-conditioned supernatants of WT, ΔAcpB and mFabT strains. A) Adhesion to endometrial cells; B) Growth in endometrial cell conditioned supernatant. Growth experiments were started with 10_3_ bacteria per ml. Bars and symbols, white, WT strain; blue, ΔAcpB strain; green, mFabT strain. N= 8; one-way ANOVA, Bonferroni post-test, *, p<0.05, **, p<0.001.

In contrast, to the mFabT strain, no growth defect was observed between the WT and the ΔAcpB strain (Fig. 3B) (7). Since mFabT growth defect was attributed to energy loss due to unrepressed FA synthesis, this suggests that the slight FASII gene repression observed in the ΔAcpB strain in the presence of eFA is sufficient to restrict this energy loss, thus to impede its deleterious consequence.

In conclusion, AcpB is FabT main co-repressor in *S. pyogenes* and most probably other streptococci. This is in agreement with the increased DNA-binding affinity provided by AcpB but not by AcpA in *S. pneumoniae* in *in vitro* conditions (8-10). That a deleted *S. pyogenes acpB* strain grows like the WT strain in laboratory media whereas deleted *fabT* strain could not be obtained or could barely grow indicates a functional redundancy that is provided by AcpA, supporting that AcpB is not, in *S. pyogenes*, the sole corepressor (7, 13, 14). In other species, such as *Streptococcus agalactiae* where *acpB* is absent or severely truncated in over half the sequenced strains and *E. faecalis* where transcription repression is partially retained in a Δ*acpB* strain, AcpB has weaker role (5-8). Finally, in *S. pyogenes* AcpB deletion weakens the bacteria - host cell interaction supporting the importance of the FASII regulation for bacterial interaction with its environment.

## Material and Methods

### Bacterial strains and culture conditions

The strains used in this study are described in Table S1. *S. pyogenes* strains were grown under static condition at 37 °C in Todd Hewitt broth supplemented with 0.2 % Yeast Extract (THY) or on THY agar plates (THYA). To study, the role of eFA addition or eFA incorporation, the medium was supplemented with 0.1 % Tween 80 (THY-Tween) (Sigma-Aldrich, P1754), essentially composed of C18:1Δ9 or 100 μM C17:1 (Larodan, Sweden). Overnight cultures were diluted to an OD_600nm_ = 0.05 and grown in the indicated medium to the exponential phase (OD_600nm_ comprised between 0.4 and 0.5).

### Strain construction

The primers used for the generation of the plasmids and verifying the different strains are described in Table S3. The ΔAcpB were obtained by homologous recombination of the plasmid pG1AcpB following the same protocol as described previously (15, 16). The DNA fragments encompassing *fabT* were cloned in BamHI – EcoRI digested pG1 using the In Fusion cloning kit® (Clonetech) (17). This led to the deletion of *acpB* from nucleotides 14 to 203, relative to the translation initiation site. The back-to-the wild-type (BWT), *i*.*e*. reversion of the single cross-over and restoration of a wild-type *acpB*, was carried out as previously described. The whole genomes of the constructed strains were sequenced; one identical surreptitious mutation was found in both the ΔAcpB (Bioproject PRJNA926803, accession number, SAMN34258008) and the BWT (accession number, SAMN34258009) strains in *priA*, the G in position 1073 was replaced by a T leading to the replacement of the 358^th^ amino acid residue, a tryptophane by a leucine.

### RNA isolation and First-strand cDNA synthesis, quantitative PCR (qPCR)

*S. pyogenes* strains were cultured at 37 °C in THY or THY-Tween, and cells were harvested at exponential growth phase (OD_600nm_ comprised between 0.4 and 0.5). RNA isolation, cDNA synthesis and quantitative PCR were carried out as previously described (7).

### Fatty acid analysis

Strains were grown in THY, THY-Tween, or THY-C17:1 until OD_600nm_ = 0.4 - 0.5. FAs were extracted and analyzed as previously described (18))(10, 18-20) . Briefly, analyses were performed in a split-splitless injection mode on an AutoSystem XL Gas Chromatograph (Perkin-Elmer) equipped with a ZB-Wax capillary column (30 m x 0.25 mm x 0.25 mm; Phenomenex, France). Data were recorded and analyzed by TotalChrom Workstation (Perkin-Elmer). FA peaks were detected between 12 and 40 min of elution, and identified by comparing to retention times of purified esterified FA standards (Mixture ME100, Larodan, Sweden). Results are shown as percent of the specific FA compared to total peak areas (TotalChrom Workstation; Perkin Elmer).

### Cell culture

HEC-1-A (ATCC_ HTB-112TM) endometrial epithelial cells were cultured as recommended, in McCoy’s 5A medium (Gibco, Ref. 26600080) supplemented with 10 % fetal bovine serum at 37 °C, 5 % of CO_2_.

### Bacterial adhesion capacity

*S. pyogenes* adhesion capacity was realized as described previously after growing the bacteria in THY to an OD_600nm_ of 0.4 to 0.5; bacteria were added at a multiplicity of infection of 1 (15). The values were normalized to the inoculum for each experiment.

### Bacterial growth capacity in the presence of eukaryotic cells or their culture supernatant

*S. pyogenes* bacteria were grown in THY to an OD_600nm_ of 0.4 to 0.5, the culture was washed twice in PBS and diluted in RPMI medium without glutamine (Gibco, Ref. 32404-014) to inoculate filtered cell culture supernatants (conditioned supernatants) at a final concentration of 10^3^ bacteria per ml as previously described (7). After 8 h incubation, serial dilutions were plated on THY-agar plates. The number of CFUs was determined after 24 h of growth at 37 °C and normalized to the inoculum for each experiment.

### Statistical analysis

Data were analyzed with GraphPad Prism version 9.4.1. The tests used are indicated in the figure legends. Statistical significance is indicated by: ns (not significant, p > 0.05); *, p < 0.05; **, p < 0.01; ***, p < 0.001; ****, p < 0.0001.

## Acknowledgements

We thank A. Gruss for her constant interest and support. We thank Tiphaine Pouillon and Maëva Gauduin undergraduates in the laboratory for technical help.

CL was supported by Université Paris Cité (BioSPC, n°51809666), FRM (FDT202106012831) and FEMS (FEMS Congress Attendance Grant for poster n° 7875 in 2021 and FEMS grant - LISSSD 2022 n° LISS-213065). QZ was supported by bourse SSHN jeune chercheur and innovative research team of high-level local universities in Shanghai (SHSMU-ZDCX20212700). This work was supported by DIM One Health (RPH17043DJA).

