## Supplementary Figures 1 to 4 for "Acyl-AcpB, a FabT co-repressor in *Streptococcus pyogenes*"

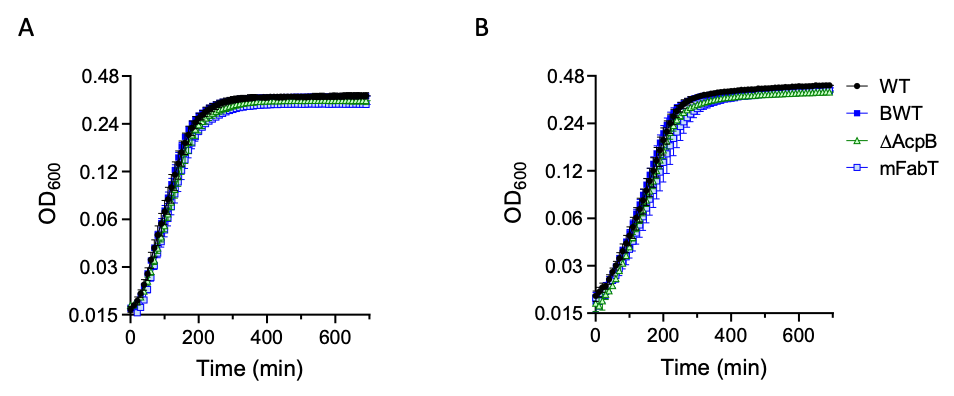

**Figure S1. Impact of *acpB* deletion on *S. pyogenes* growth capacity in rich media.** Growth curves of WT, mFabT, AcpB and BWT strains, in A) THY and B) THY-Tween media. Growth was monitored in a Multiscan (Thermo Scientific).

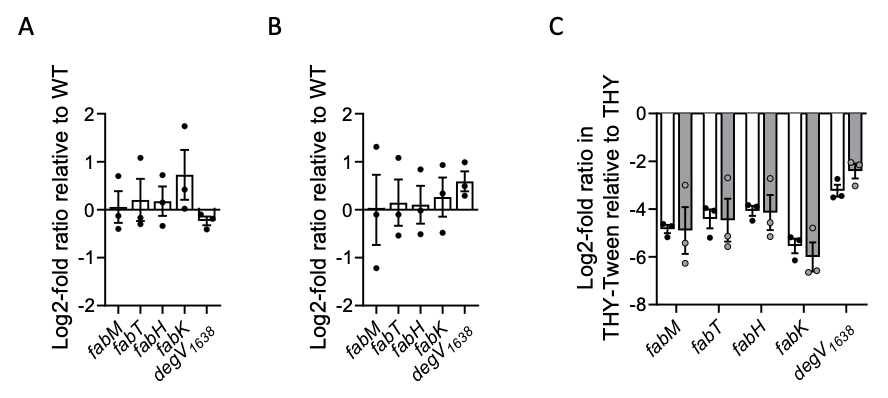

**Figure S2. Comparison of FASII and the FabT-regulated *degV_1638_* gene repression in *S. pyogenes* WT and BWT strains.** Strains were grown in THY (A, C) or THY-Tween 80 (B, C) and RNAs were quantified by qRT-PCR. Expression was normalized to that of *gyrA*; (A, B) relative gene expression is expressed as the log2-fold ratio in BWT *versus* WT strain. C) The relative gene expression is expressed as the log2-fold ratio between a given strain grown in THY Tween 80 *versus* THY, white bars, WT; grey bars, BWT

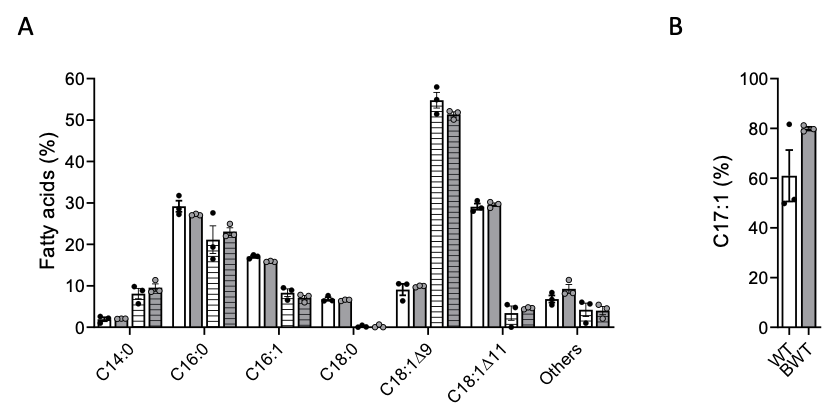

**Figure S3. FA membrane composition of WT and BWT strains.** Quantified proportions of major FAs of WT and BWT strains grown in A) THY or THY-Tween 80, B) THY-17:1; bars, white, WT strain; grey, BWT strain; A) open, THY; hatched, THY-Tween 80. Comparisons were carried out between WT and BWT strains; none were statistically different.

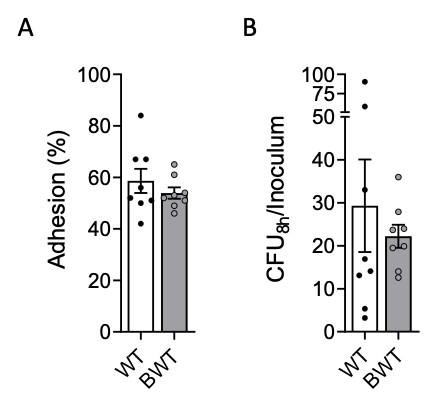

**Figure S4. Comparison of the BWT and WT adhesion and growth capacity**. A) Adhesion to human endometrial cells and B) growth capacities in endometrial cell conditioned supernatants of WT and BWT strains. Growth experiments were started with 10^3^ bacteria per ml. Bars and symbols, white, WT strain; grey, BWT strain N= 8; T-test, none were statistically different.
