## Supplementary methods for "Acyl-AcpB, a FabT co-repressor in *Streptococcus pyogenes*"

**Growth curves.** *S. pyogenes* stationary precultures were diluted in THY or THY-Tween 80 to an OD_600nm_ = 0.05 and transferred to 96-well plates. These were incubated at 37 °C in a Thermo Scientific Multiskan GO (ThermoFischer Scientific) and growth was determined by measuring absorbance at 600 nm every 10 min after shaking the plates.
